# Neonatal enteroids absorb extracellular vesicles from human milk-fed infant digestive fluid

**DOI:** 10.1101/2023.09.03.556067

**Authors:** Claire Yung, Yang Zhang, Madeline Kuhn, Randall J. Armstrong, Amy Olyaei, Molly Aloia, Brian Scottoline, Sarah F. Andres

## Abstract

Human milk contains extracellular vesicles (HMEVs). Pre-clinical models suggest that HMEVs may enhance intestinal function and limit inflammation; however, it is unknown if HMEVs or their cargo survive neonatal human digestion. This limits the ability to leverage HMEV cargo as additives to infant nutrition or as therapeutics. This study aimed to develop an EV isolation pipeline from small volumes of human milk and neonatal intestinal contents after milk feeding (digesta) to address the hypothesis that HMEVs survive in vivo neonatal digestion to be taken up intestinal epithelial cells (IECs). Digesta was collected from nasoduodenal sampling tubes or ostomies. EVs were isolated from raw and pasteurized human milk and digesta by density-gradient ultracentrifugation following two-step skimming, acid precipitation of caseins, and multi-step filtration. EVs were validated by electron microscopy, western blotting, nanoparticle tracking analysis, resistive pulse sensing, and super-resolution microscopy. EV uptake was tested in human neonatal enteroids. HMEVs and digesta EVs (dEVs) show typical EV morphology and are enriched in CD81 and CD9, but depleted of β-casein and lactalbumin. HMEV and some dEV fractions contain mammary gland-derived protein BTN1A1. Neonatal human enteroids rapidly take up dEVs in part via clathrin-mediated endocytosis. Our data suggest that EVs can be isolated from digestive fluid and that these dEVs can be absorbed by IECs.

## 1. INTRODUCTION

Human milk (HM) contains numerous bioactive factors with nutritive and nonnutritive benefits to promote the health and long-term well-being of the developing infant (1). For example, feeding human milk enhances intestinal function and reduces the risk of necrotizing enterocolitis (2–10), promotes healthy brain development (11), and reduces the risk of chronic disorders, such as asthma or inflammatory bowel disease (12).

Human milk is replete with extracellular vesicles (EVs), which are nanoparticle-sized structures secreted by most cells within the human body (13). EVs, including human milk EVs (HMEVs), carry biological information between donor and recipient cells, to confer changes in cellular function or biological activity (14–22). HMEVs have putative beneficial effects, as shown by a combination of in vitro and animal studies demonstrating that HMEVs promote intestinal epithelial cell (IEC) proliferation (16, 19, 20), reduce experimental necrotizing enterocolitis (NEC), limit inflammatory (15–17, 21, 22) or non-inflammatory damage (15, 19, 23), promote expression of intestinal barrier proteins (21), or enhance epithelial barrier function (15, 18). Collectively, these data suggest that HMEVs confer functional benefits on IECs and hold immense therapeutic promise for humans. To do so, however, HMEVs must survive infant digestion to reach the intestinal epithelium.

Several in vitro studies indicate that HMEVs are minimally impacted by a pH of 4-4.5 mirroring that of a term or preterm infant stomach, as well as incubation with several gastrointestinal proteases (24–26). More recent work shows that bovine milk EVs are intact after exposure to a pH as low as 1.3 (15). Furthermore, existing animal studies indicate that bovine or murine milk-derived EVs survive murine digestion to reach the placenta and embryo (27) or intestine and brain (28, 29). Studies also demonstrate that HMEVs confer beneficial effects within the murine intestine (15, 17, 21), suggesting that they too may survive murine digestion. Yet it remains unknown if HMEVs survive neonatal human digestion to functionally impact the neonatal human intestinal epithelium. Answering this question is a critical next step for understanding the ability to orally deliver HMEVs as therapeutics or drug carriers.

The neonatal intestinal epithelium is thought to take up macromolecules through bulk endocytosis (30, 31), a mechanism unique to the neonate where endocytic cargo is broken down by the lysosome. Existing data similarly suggest that endocytosis is the primary mechanism of EV uptake, although precise mechanisms may vary depending on the cell of EV origin and the acceptor cell type (32, 33). Studies examining bovine milk EV uptake in mice or human cell lines (Caco2) indicate that the FcRn receptor plays a role in the uptake of some milk EVs (34, 35).

Studies examining HMEV uptake are limited and use transformed IEC lines, such as Caco2 (24) or HIEC (25) cells. Although these cell lines are commonly used to model the small intestinal epithelium (36, 37), Caco2 cells are derived from human colorectal cancer (38, 39), while HIEC cells are derived from 17-19 week fetal ileum (40). To date, the uptake of HMEVs by primary, human neonatal IECs remains unexplored.

In this study, we hypothesized that a proportion of HMEVs in human milk survive in vivo infant digestion to reach the small intestine and be taken up by IECs. To test this hypothesis, we utilized a rare and valuable set of fed human milk and intestinal contents (digesta) samples isolated from infants in the neonatal intensive care unit (NICU). We optimized a relatively high-purity EV isolation pipeline allowing for the isolation of HMEVs from 1 mL of raw human milk (RHM), pasteurized donor human milk (PDHM), or digesta. We also developed and characterized a primary human neonatal enteroid culture system to examine digesta EV (dEV) uptake ex vivo. Enteroids are 3D organoid cultures grown from intestinal stem cells (41, 42). In contrast to immortalized cell lines used in previous studies (24, 25), primary neonatal enteroids are arguably a more physiologically relevant model for studying normal IEC function (43) as they are derived from normal infant intestinal stem cells and develop multiple differentiated IEC types.

Our data demonstrate for the first time that EVs can be isolated from human milk-fed infant digesta. These dEVs are readily and robustly endocytosed by neonatal enteroids containing differentiated IECs, primarily via dynamin-dependent processes. This study addresses a critical gap in understanding the potential role(s) of HMEVs in the infant intestine and the pipeline for HMEV therapeutic development.

## 2. MATERIALS AND METHODS

All human studies were approved by the Oregon Health & Science University (OHSU) Institutional Review Board (IRB) IRB #17968 and #21952. Milk and digesta samples were collected after obtaining parental informed consent; tissue was obtained via IRB-approved processes.

### 2.1 Sample collection

Human milk samples, collection, and storage are listed in **Tables 1** and **2**. All samples were aliquoted and frozen before processing and every effort was taken to minimize freeze thaws. Raw human milk (RHM) or pasteurized donor human milk (PDHM) collection was performed under OHSU IRB #17968. PDHM was obtained from the Northwest Mother’s Milk Bank (Beaverton, OR, USA) or Prolacta (City of Industry, CA, USA).

**Table 1.**
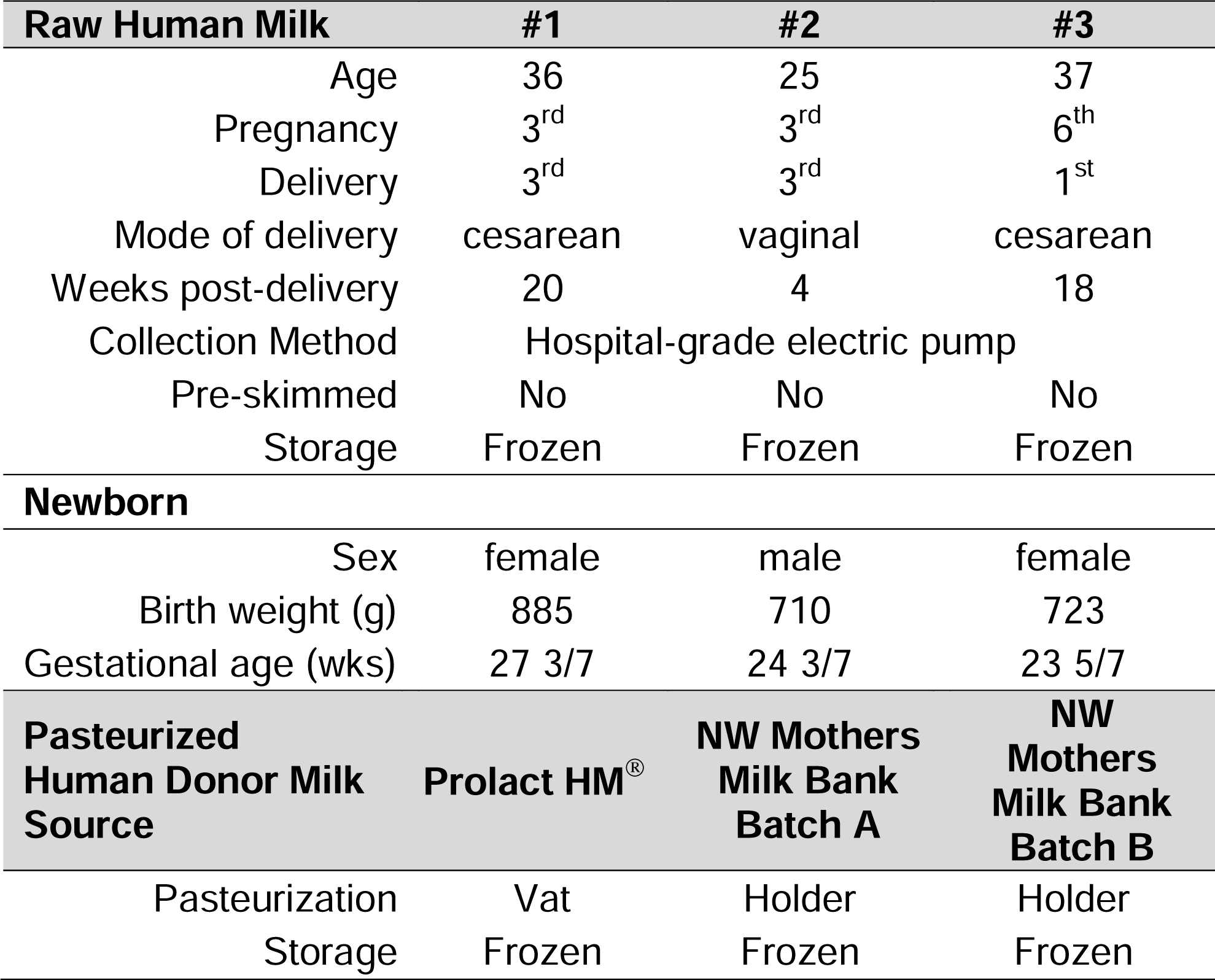
Raw and pasteurized human milk sample demographics.

| Raw Human Milk | #1 | #2 | #3 |
| --- | --- | --- | --- |
| Age | 36 | 25 | 37 |
| Pregnancy | 3<sup>rd</sup> | 3<sup>rd</sup> | 6<sup>th</sup> |
| Delivery | 3<sup>rd</sup> | 3<sup>rd</sup> | 1<sup>st</sup> |
| Mode of delivery | cesarean | vaginal | cesarean |
| Weeks post-delivery | 20 | 4 | 18 |
| Collection Method | Hospital-grade electric pump |  |  |
| Pre-skimmed | No | No | No |
| Storage | Frozen | Frozen | Frozen |
| **Newborn** |  |  |  |
| Sex | female | male | female |
| Birth weight (g) | 885 | 710 | 723 |
| Gestational age (wks) | 27 3/7 | 24 3/7 | 23 5/7 |
| Pasteurized Human Donor Milk Source | Prolact HM<sup>®</sup> | NW Mothers Milk Bank Batch A | NW Mothers Milk Bank Batch B |
| Pasteurization | Vat | Holder | Holder |
| Storage | Frozen | Frozen | Frozen |

**Table 2.** Matched human milk and digesta sample demographics.

| Raw Human Milk | Milk (a) | Milk (b) | Milk (c) |
| --- | --- | --- | --- |
| Age | 30 | 36 | 29 |
| Pregnancy | 3rd | 1st | 1st |
| Delivery | 3rd | 1st | 1st |
| Mode of delivery | cesarian | cesarian | cesarian |
| Weeks post-delivery | 7 | 11 | 6 |
| Collection Method | Hospital-grade electric pump |  |  |
| Pre-skimmed | No | No | Yes |
| Storage | Frozen | Frozen | Frozen |
| Digesta (d) | d(a) | d(c) | d(c) |
| Feed | Raw HM | Raw HM | Raw HM |
| Sex | Male | Male | Male |
| Gestational age (wks) | 44 4/7 | 44 1/7 | 40 3/7 |
| Collection Location | 2<sup>nd</sup> duo | 4th duo | jejunum |
| Collection Method | PPT | PPT | Ostomy |
| Pre-skimmed | No | No | Yes |
| Sample Storage | Frozen | Frozen | Frozen |

PPT = post-pyloric collection tube

In vivo, digested milk samples (digesta) were collected from infants in the Doernbecher Children’s Hospital NICU at OHSU under IRB #17968. Infant inclusion criteria were: infants already admitted to the NICU, greater than 26 weeks post-menstrual age, presence of an indwelling nasogastric or orogastric feeding tube, and tolerating full enteral feeding volumes. Infants were excluded from the study if they had anatomic or functional gastrointestinal disorders that would affect digestion, were medically unstable, or were non-viable. Digested milk samples were collected from three infants (**Table 2**). Feeds were delivered via a gastric tube over 30 min or less. For samples (a) and (b), a post-pyloric collection tube (PPT) was placed into the duodenum or proximal jejunum before feeding. Samples were collected via gravity flow as the digesta passed the PPT port and dripped into sterile vials, placed on ice. The samples were aliquoted and stored at –80 °C. Sample (c) was collected from a fresh, sterile ostomy bag as the digesta exited the stoma. Stoma output was only allowed to accumulate in the bag long enough to reach a feasibly collected volume.

### 2.2 Isolation of extracellular vesicles from human milk and neonatal digesta

Samples were quickly thawed in a 37 °C water bath and used immediately (44, 45). A step-by-step illustration is depicted in **Figure 1**. One milliliter of RHM, PDHM, or digesta was used for each isolation. Samples were centrifuged at 6,000 x g for 10 min at 4 °C to remove cream and large cellular debris (46). The supernatant was carefully collected and centrifuged again at 2,000 x g for 20 min at 4 °C to further remove any fat globules and/or smaller cellular debris. This supernatant was then passed through Whatman^TM^ Grade 1 filter paper (diameter 42.5 mm) using a glass funnel before isoelectric casein precipitation.

**Figure 1.**
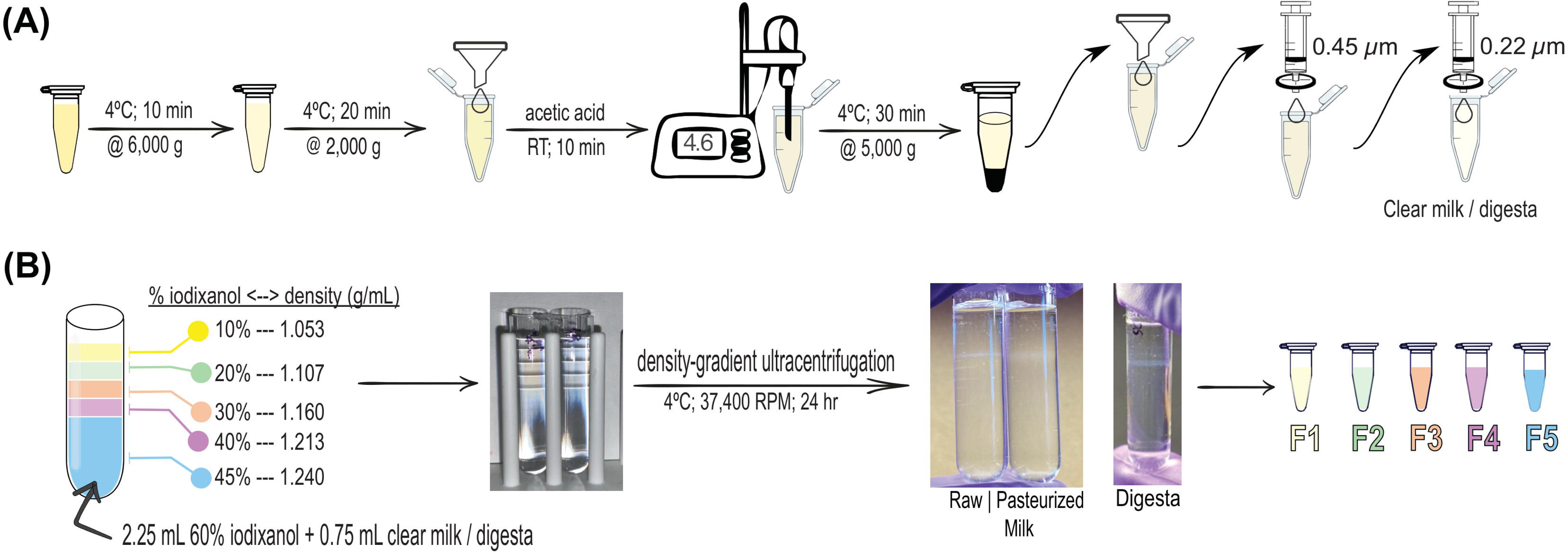
Human milk (HM) and neonatal digesta extracellular vesicle (dEV) isolation pipeline. (A) Samples were quickly thawed in a 37 °C water bath and used immediately (44, 45). One milliliter of whole HM was used for each isolation. Samples were centrifuged at 6,000 x g for 10 min at 4 °C to remove cream and large cell debris (46). The supernatant was carefully collected and centrifuged again at 2,000 x g for 20 min at 4 °C to further remove any fat globules and/or smaller cell debris. This supernatant was then passed through Whatman™ Grade 1 filter paper (diameter 42.5mm) using a glass funnel before isoelectric casein precipitation. Acetic acid (1 and 10 N) was used to adjust the pH of the supernatant to 4.6 using a portable pH meter. pH-adjusted samples were incubated at room temperature for 10 min. Casein was pelleted by centrifugation at 5,000 x g for 30 min at 4 °C. The supernatant was collected and passed sequentially through Whatman™ Grade 1 filter paper (diameter 42.5 mm) using a glass funnel, a 0.45 µm syringe filter, and a 0.22 µm syringe filter for casein removal. (B) Bottom-up density gradient ultracentrifugation was performed using iodixanol gradient (47). A 45% (w/v) bottom layer contained OptiPrep™ solution with clear HM or digesta (2.25 mL OptiPrep™ + 0.75 mL sample = 3 mL) into a 5 mL open-top thin wall ultra-clear tube. The discontinuous gradient with 40% (w/v), 30% (w/v), 20% (w/v), and 10% (w/v) solutions of iodixanol (diluted with sterile, 0.1 µm-filtered PBS) were carefully layered on top, drop by drop. Samples were spun in an ultracentrifuge at 37,400 rpm for 24 hrs at 4 °C with zero acceleration and deceleration. After ultracentrifugation, 500 µL fractions were carefully collected into individual Protein LoBind™ (Eppendorf) tubes from top to bottom using a P1000 pipettor without disrupting the layers. The fractions were used immediately for downstream analysis.

Acetic acid (1 and 10 N) was used to adjust the pH of the supernatant (initial pH 6.8 - 7.4) to 4.6 using a portable pH meter (Southern Labware, Fisher Scientific, Hampton, NH, USA). Acetic acid was added to samples ≤1 µL at a time, and samples were briefly mixed after each acid addition. Once the pH was adjusted to 4.6, samples were incubated at room temperature for 10 min before final centrifugation at 5,000 x g for 30 min at 4 °C. The supernatant was carefully collected and was passed through Whatman^TM^ Grade 1 filter paper (diameter 42.5 mm) using a glass funnel, a 0.45 µm syringe filter, and a 0.22 µm syringe filter to remove casein. The residual fluid yielded clarified HM or digesta.

Density gradient ultracentrifugation using a SW Type 55 Ti swinging bucket rotor in LM-70 ultracentrifuge (Beckman Coulter, Brea, CA, USA) was achieved using OptiPrep^TM^ iodixanol solution (60% iodixanol, w/v) following the bottom-up technique (47). A 45% (w/v) bottom layer was made by diluting OptiPrep™ solution with clarified HM or digesta sample (2.25 mL OptiPrep™ + 0.75 mL sample = 3 mL) into a 5 mL open-top thin wall ultra-clear tube (Beckman Coulter). A discontinuous gradient was made with 40% (w/v), 30% (w/v), 20% (w/v) and 10% (w/v) solutions of iodixanol (diluted with sterile, 0.1 µm-filtered phosphate-buffered saline (PBS)) carefully layered on top of the prior density, drop by drop. The ultracentrifugation was carried out at 169,522 × g (37,400 rpm) for 24 hours at 4 °C with zero acceleration and deceleration. After ultracentrifugation, 500 µL fractions were carefully collected into protein LoBind tubes (Fisher) from top to bottom using a P1000 pipette without disrupting the layers. The fractions were used the same day for analysis or functional studies or else stored at 4 °C. We submitted all relevant data of our experiments to the EV-TRACK knowledgebase (EV-TRACK ID: EV230590) (48).

### 2.3 Cell Lines

SW480 and MCF7 cells (western blot positive controls) were cultured in DMEM high glucose medium (Gibco, Thermo Fisher, Waltham, MA, USA) supplemented with 10% fetal bovine serum (FBS, Cytiva HyClone™, Marlborough, MA, USA) and 1% penicillin/streptomycin (Gibco), and grown at 37 °C in a humidified incubator with 5% CO_2_. Cells were lysed in radioimmunoprecipitation assay-sodium dodecyl sulfate (RIPA-SDS) and protein concentration was quantified using a bicinchoninic acid (BCA) reagent kit (Thermo Fisher, Waltham, MA, USA) following manufacturer instructions ahead of western blotting.

### 2.4 Validation of extracellular vesicle isolation

#### 2.4.1 Western blotting

For validation, EV preparations and cellular controls were lysed in 6X Laemmli buffer (375 mM Tris-HCl pH 6.8, 9% SDS w/v, 50% Glycerol v/v, 0.03% Bromophenol blue w/v, 9% DTT w/v), incubated at 37 °C for 30 min, and loaded onto 4 to 12% Bis-Tris mini protein gels (NuPAGE™, Invitrogen, Waltham, MA, USA, 1.5 mm). PageRuler Plus™ Prestained Protein Ladder (Thermo Fisher, 10 – 250 kDa) was resolved as a molecular weight reference. Human cell lines, MCF7 (breast cancer) (ATCC) and SW480 (colorectal cancer) (ATCC) lysates were also resolved as cellular controls. The ATCC authenticates all human cell lines through short tandem repeat analysis. All cell lines were confirmed to be mycoplasma negative every 3 weeks (MycoAlert™ Mycoplasma Detection Kit, Lonza, Bend, OR, USA). Fifteen micrograms of total whole cell lysates (WCL) and 20 µL of EV lysate isolated from 1 mL starting volume were resolved by gel electrophoresis. The samples were stacked for 30 min at 60V and resolved for 60 min at 150V in 1X NuPAGE™ MOPS SDS running buffer (Invitrogen) and transferred onto 0.45µm PVDF membrane (Immobilon, Millipore/Merck) via a wet transfer system with 1X NuPAGE^TM^ transfer buffer (Invitrogen) at 100 V for 60 min. Membranes were blocked with LI-COR PBS blocking buffer (LI-COR, Lincoln, NE) at room temperature for 60 min.

MISEV 2018 guidelines (13) were used to characterize the presence of EV-specific proteins via the following primary antibodies: 1) anti-transmembrane protein markers: anti-CD9 (rabbit, Abcam, Cambridge, UK, ab236630 1:1000); anti-CD63 (rabbit, System Biosciences, Palo Alto, CA, USA, EXOAB-CD63A-1, 1:1000); anti-CD81 (mouse, Santa Cruz Biotechnology, Santa Cruz, CA, USA, sc-166029, 1:1000), and 2) cytosolic protein markers: anti-TSG101 (mouse, BD Biosciences, Franklin Lakes, NJ, USA, 612697, 1:1000) and anti-FLOT1 (rabbit, Cell Signaling Technology, Danvers, CT, USA, 3253S, 1:10,000-1:15,000). Lactating mammary gland protein butyrophilin (BTN1A1) anti-BTN1A1 (mouse, OriGene Technologies, Rockville, MD, USA, TA501529S 1:3000). Anti-ITGB1, (rabbit, Cell Signaling Technology, 34971T, 1:1000) was used to identify microvesicle contamination (49). Anti-GM130 (rabbit, Novus Biologicals, Centennial, CO, USA, NBP2-53420, 1:1000) was used to identify cis-Golgi matrix protein as a marker of cellular contamination (13). Anti-LALBA (rabbit, Abcam, ab178431, 1:1000) and anti-CSN2 (rabbit, Abcam, ab205301, 1:5000) assessed non-HMEV components from lactalbumin and casein, respectively. All primary antibodies were diluted in 0.5X LI-COR PBS blocking buffer (LI-COR Biosciences, Lincoln, NE, USA) diluted with 1X PBS containing 0.2% tween-20.

The membranes were incubated with goat anti-rabbit labeled with IRDye 700 (1:20 000, v/v, LI-COR) and/or goat anti-mouse antibody labeled with IRDye 800 (1:20,000, v/v, LI-COR) for 60 min before visualization on LICOR Odyssey. All secondary antibodies were diluted in 0.5X LI-COR PBS blocking buffer diluted with 1X PBS containing 0.2% tween-20 and 0.01% sodium dodecyl sulfate (SDS).

#### 2.4.2 Nanoparticle Tracking Analysis

All samples were diluted in 10% particle-free PBS (Genesee Scientific, Morrisville, NC, USA) to a final volume of 1 mL. Samples were diluted to achieve 50 – 250 particles/frame) before injection (50). Samples were analyzed using a ZetaView^®^ Twin (Particle Metrix, Ammersee, Germany). Each sample was run in triplicate by scanning 11 cell positions under the following settings: Focus: autofocus; Frame rate: 30; Camera sensitivity for all samples: 70 for scatter, 87 for Di8; Shutter: 100; Scattering Intensity: detected automatically; Cell temperature: 25 °C. The videos were analyzed with the built-in ZetaView^®^ Software 8.05.12 SP2 with specific analysis parameters: Maximum area: 1000, Minimum area 10, Minimum brightness: 20. Hardware: embedded laser: 488 and 520 nm. Di-8-ANEPPS, final concentration 2 µM, (Thermo Fisher Scientific) was used to stain HMEVs as described (51); camera: CMOS. EV concentration (EV particles/mL) was calculated by accounting for dilution factors.

The characteristics of fractions 2 and 3 were further validated by treating Di8-stained samples with 0.1% Triton-X100 (final concentration), 5 mM EDTA (final concentration), 50 µg/mL proteinase K (Thermo Fisher Scientific, final concentration), or 0.1% Triton-X100 + 50 µg/mL proteinase K. Samples were incubated for 5-10 minutes at room temperature and particle counts were measured on the Zetaview^®^ using the same parameters as above. Milk sample (c) was used for the experiment.

#### 2.4.3 Resistive Pulse Sensing (RPS)

HMEV and dEV samples were diluted 1:20 in 1% Tween-20 (v/v) in 0.2µm filtered 1X PBS (diluent). Samples were loaded into polydimethylsiloxane cartridges (diameter range 65 nm to 400 nm, C-400) with mold ID 108H. A new cartridge was used for each sample. Measurements were collected using a Spectradyne nCS1™ instrument (Spectradyne, Signal Hill, CA, USA). The instrument was primed with a cleaning cartridge prior to use. Four thousand events were collected from 4 µL of each sample. Analysis was performed using the accompanying nCS1™ Data Viewer (Version 2.5.0.325). All data was background corrected and peak filtering was applied using the mold ID and a diameter of 80-250nm. Size and concentration measurements were calibrated based on 150 nm polystyrene beads diluted 1:10,000 in diluent. Concentrations were adjusted using Microsoft Excel™ and graphs were made in GraphPad Prism^®^ 10 (GraphPad Software, San Diego, CA, USA).

#### 2.4.4 Morphological characterization of extracellular vesicles by negative transmission electron microscopy (TEM)

Electron microscopy was performed at the Multiscale Microscopy Core, an Oregon Health & Science University Shared Resource facility. Briefly, EV preparations were fixed in s 4% paraformaldehyde (PFA) solution (Thermo Scientific). Five microliters of fixed EVs were deposited onto glow discharged (60 secs 15 mA, negative mode) carbon formvar 400 Mesh copper grids (Ted Pella 01822-F) for 3 min, rinsed 30 sec in water, wicked on Whatman filter paper 1, stained for 60 sec in filtered 1.33% (w/v) uranyl acetate in water, wicked and air dried. Samples were imaged at 120 kV on a FEI Tecnai™ Spirit TEM system. Images were acquired using the AMT software interface on a NanoSprint™12S-B CMOS camera system.

#### 2.4.5 Micro BCA^TM^ for protein concentration

Protein concentrations from fractions 2 and 3 of fed human milk and digesta were determined using a Micro BCA^TM^ kit (Pierce, Thermo Fisher), following manufacturer instructions.

#### 2.4.6 Super-resolution microscopy

EV surface proteins were observed and quantified using the EV Profiler Kit (EV-MAN 1.0, ONI, Oxford, United Kingdom) and direct stochastic optical reconstruction microscopy (dSTORM). EV samples were affinity captured on microfluidic chips using CD9/CD81/CD63 antibodies provided in the kit. HCT116 and MCF7 EVs were used as controls. Immobilized EVs were fixed with F1 solution (provided in the kit) for 10 minutes before labeling with CD9-CF488 (excitation (ex)/emission (em): 490/515 nm) and CD63-CF568 (ex/ em: 562/583 nm) antibodies provided in the kit. Labeled EVs were again fixed with F1 for 10 minutes. EV samples were imaged on the Nanoimager S Mark II microscope (ONI) with 100X oil-immersion objective, and labeled proteins were imaged sequentially at 35 and 50% power for the 561 and 488 nm lasers, respectively, at 1,000 frames per channel with the angle of illumination set to 52.5°. dSTORM-imaging buffer was freshly prepared and added just before image acquisition. The system was calibrated using the bead slide manual assembly (ONI) before use. Data were processed on NimOS software (version 1.19, ONI). Subpopulation analyses of EVs that express one or two markers were analyzed using ONI’s online platform CODI (https://alto.codi.bio). We used a density-based clustering analysis with drift correction and filtering to evaluate each vesicle. At least 5 localizations were required to constitute a vesicle and at least 3 localizations of one protein were required to consider the localizations a real signal.

### 2.5 Enteroid cultures

#### 2.5.1 Neonatal human enteroid generation and culture

Tissue collected for enteroid generation was approved under OHSU IRB # 21952. Enteroids were generated from small intestinal tissue isolated from the ileum of a 2-month-old participant born at 39 weeks undergoing surgery for ileal atresia. After collection, tissue was washed in sterile PBS with 1% penicillin/streptomycin and frozen in freezing media (10% FBS; 10% DMSO; Dulbecco’s modified Eagle medium/F12). Tissue was rapidly thawed in a 37 °C water bath and then washed three times in sterile PBS with 10% FBS. Luminal pinch biopsies were collected. The biopsies were minced in sterile PBS with 10% FBS using sterile scissors and spun at 300 x g for 3 min at 4 °C. The pellet was resuspended in 500 µl of “working solution”: Liberase™ TH 0.1 WU/mL (Sigma, St. Louis, MO, USA); DNAse1 50 U/mL (Roche, Indianapolis, IN, USA); Hank’s balanced salt solution (HBSS) and incubated in a Thermomixer at 800rpm for 10 min at 37 °C. During the incubation, a 100 µm strainer was washed with 8 mL of 4% bovine serum albumin (BSA) in HBSS to prevent cells from sticking to the filter. The tissue was pelleted at 300 x g for 3 min at 4 °C and the supernatant was passed through the prepared 100 µm strainer. The remaining tissue fragments were resuspended in another 500 µl of working solution and mixed in the Thermomixer^®^ at 800 rpm for 10 min at 37 °C. After incubation, the sample was passed through a P1000 pipettor tip 10 times and spun down at 300 x g for 3 min at 4 °C. The supernatant was filtered through the same 100 µm strainer and the tissue pellet was resuspended in 500 µl of working solution and mixed in the Thermomixer at 800 rpm for 10 min at 37 °C. The remaining tissue was passed through a P200 pipettor tip on the end of a P1000 tip 10 times. Once the tissue was small enough, the slurry was filtered through the same 100 µm strainer. The filtered cells were pelleted at 500 x g for 5 min at 4 °C. The pellet was resuspended in 500 µl of 4% BSA in HBSS and counted using Trypan Blue staining (Gibco) and a Countess cell counter (Invitrogen). The cells were seeded at a density of up to 10^5^ live cells per 30 µl Matrigel (Corning, Corning, NY, USA) dome (80% Matrigel). After the Matrigel solidified at 37 °C, Human Intesticult™ (StemCell Technologies, Cambridge, MA, USA) with Primocin^®^ (20 µl of a 500X stock, InvivoGen, San Diego, CA, USA) was overlaid. Primocin^®^ was discontinued after the second passage and the enteroids were supplemented with Anti/Anti (Gibco). Enteroids were maintained and expanded in Matrigel patties in the “basal out” orientation.

#### 2.5.2 Apical-out enteroids

To access the apical/luminal surface of the human enteroids, we reversed enteroid polarity using the technique described previously (52, 53). Briefly, 1 mL of ice-cold cell recovery solution (Corning) was added directly to the exposed Matrigel dome to break up the Matrigel patty. The enteroid/Matrigel mixture was transferred to a 15 mL conical tube and rotated at 4 °C for 1 hr. The enteroids were then pelleted at 300 x g for 3 min at 4 °C and washed once with advanced DMEM/F-12 basal media (Gibco). The enteroid pellet was resuspended in the desired volume of IntestiCult complete medium (Stemcell Technologies) and cultures were maintained in a 24-well ultra-low-attachment plate for 48 hours at 37 °C in a 5% CO_2_ incubator.

### 2.6 RNA isolation and qRT-PCR

To isolate RNA from enteroids, the media was removed and cell recovery solution was added to remove Matrigel. The enteroids were pelleted, washed with PBS, and pelleted again. RNA was isolated using GeneJET™ RNA Purification Kit (Thermo Scientific) or RNAqueous™-Micro Total RNA Isolation Kit (Invitrogen). High-capacity cDNA Reverse Transcription Kit (Applied Biosystems™, Thermo Fisher) was used to synthesize cDNA.

The following primer-probe sets were used: leucine-rich repeat containing G protein (*LGR5*, Hs00969422_m1), a marker of proliferation Ki-67 (*Ki67*, Hs00606991_m1), proliferating cell nuclear antigen (*PCNA*, Hs00696862_m1), polycomb complex protein BMI1 proto-oncogene (*BMI1*, Hs00180411_m1), DEF6 guanine nucleotide exchange factor (*DEF6*, Hs00427001_m1), intracellular adhesion molecule 2 (*ICAM2*, Hs00609563), bestrophin 4 (*BEST4*, Hs00396114_m1), sucrase-isomaltase (*SI*, Hs00356112_m1), chromogranin A (*CHGA*, Hs00900370_m1), mucin 2 (*MUC2*, Hs00159374_m1).

### 2.7 EV labeling with CMTPX

Five hundred microliters dEVs were stained with 4 µL 1 mM CMTPX dye (Invitrogen) in a 37° C water bath for 1 hour in the dark. The excess dye was removed using Amicon^®^ Ultra-0.5 100kDa MWCO centrifugal filter (Millipore, Burlington, MA, USA), and dEVs were concentrated to the desired volume for uptake experiments.

### 2.8 Uptake of dEVs by human neonatal enteroids

CMTPX-labeled dEVs (40 µL) were added to apical-out enteroids and incubated for 1.5 hours in the following conditions: at 37 °C with or without 200 µM Dynasore (Abcam), or at 4 °C. After dEV treatment, enteroids were pelleted and washed one time with 1X PBS and fixed in 4% (w/v) PFA (Fisher Scientific) for 15 min at room temperature. Fixed enteroids were pelleted and then incubated with 0.1% Triton-X for 15 min at room temperature, pelleted, and incubated with 165 nM Phalloidin (Fisher Scientific) for 40 min, followed by 600 nM DAPI (Thermo Fisher) for 10 min at RT in the dark. Enteroids were pelleted and mounted using Molecular Probes Prolong™ Gold Antifade Mountant (Thermo Fisher) and sealed with CoverGrip Coverslip sealant (Biotium, Fremont, CA, USA).

### 2.9 Confocal microscopy and image processing

dEV uptake was observed using a Zeiss LSM980 confocal laser microscope with a 20X, 0.8 NA objective (Carl Zeiss Microscopy GmbH), and lasers set at 405 nm (DAPI), 488 (Phalloidin), and 555Dnm (CMTPX) wavelengths. Images were acquired using 4X line averaging using GaAsP PMT and multi-alkali PMT detectors. Z stacks were taken through the whole enteroid (36.33 - 77.6µm). Images were processed using ZEN™ v2.5 Blue software (Carl Zeiss Microimaging GmbH). All images were acquired and processed using identical intensity settings unless noted. CMTPX uptake was quantified using ImageJ and images taken with a Keyence BZ-X800 (Keyence, Itasca, IL, USA) using a 2X objective, 0.1 NA.

### 2.10 Immunostaining

The image with immunostaining for BTN1A1 is from the Human Protein Atlas (54, 55). Lactating human mammary gland tissue was stained for BTN1A1 using Atlas Antibodies (Bromma, Sweden, Cat#HPA011126, RRID: AB_1845491; 0.1275 mg/mL; Rabbit pAb) (55).

### 2.11 Statistical analyses

Data were analyzed with GraphPad Prism^®^ 10. Diagrams are presented either as mean ± SEM and symbols represent biological replicates, either individual participant samples or enteroid passages. For the dEV uptake experiment, symbols represent individual enteroids from a single experiment.

## 3. RESULTS

This study utilized raw and pasteurized human milk, as well as matched milk digesta (neonatal intestinal contents) samples to examine whether HMEVs reach the neonatal intestine in vivo to be absorbed by IECs.

### 3.1 A pipeline to isolate HMEVs and dEVs from small sample volumes

Human milk (HM) is a complex and heterogeneous biological fluid containing, numerous proteins (including highly abundant casein and immunoglobulins), sugars, emulsified fat globules, small molecules, electrolytes, and EVs (1). Neonatal digesta is also a heterogenous biofluid containing a mixture of fed HM, stomach, and digestive secretions.

Isolating EVs from heterogeneous biofluids is challenging, especially when starting volumes are limited and relatively high purity is desired. We adapted the published protocol from Mukhopadhya et al (46) for 1 mL sample volumes followed by bottom-up, rate zone iodixanol density gradient centrifugation to isolate largely pure EVs from HM and neonatal digesta samples (**Figure 1A and B**).

### 3.2 Pipeline effectively isolates raw and pasteurized HMEVs

We validated our pipeline using raw human milk (RHM) and pasteurized donor human milk (PDHM), respectively (**Table 1**). RHM milk sample #1 and PDHM from the NW Mother’s Milk Bank Batch A were used for **Figure 2**. RHM-derived EVs were enriched in fractions (F)2 and 3, and PDHM-derived EVs were enriched in fraction 3 (**Figure 2**). Negative transmission electron microscopy (TEM) illustrated the enrichment of classic cup-shaped-EV structures ∼ 100 – 200 nm in F2 -F3 of RHM and F3 from PDHM (**Figure 2A**). Nanoparticle tracking analysis (NTA) using the lipid membrane dye Di-8-ANEPPS showed that F2-3 RHM and F3 PDHM were enriched for ∼ 200 nm-sized particles consistent with TEM data (**Figure 2B**). Based on NTA measurements of F2 and F3 for RHM and F3 of PDHM, HMEV size was not affected by pasteurization, (207.06 ± 7.86 nm and 209.68 ± 18.93 nm, n=3 RHM and PDHM, respectively).

**Figure 2.**
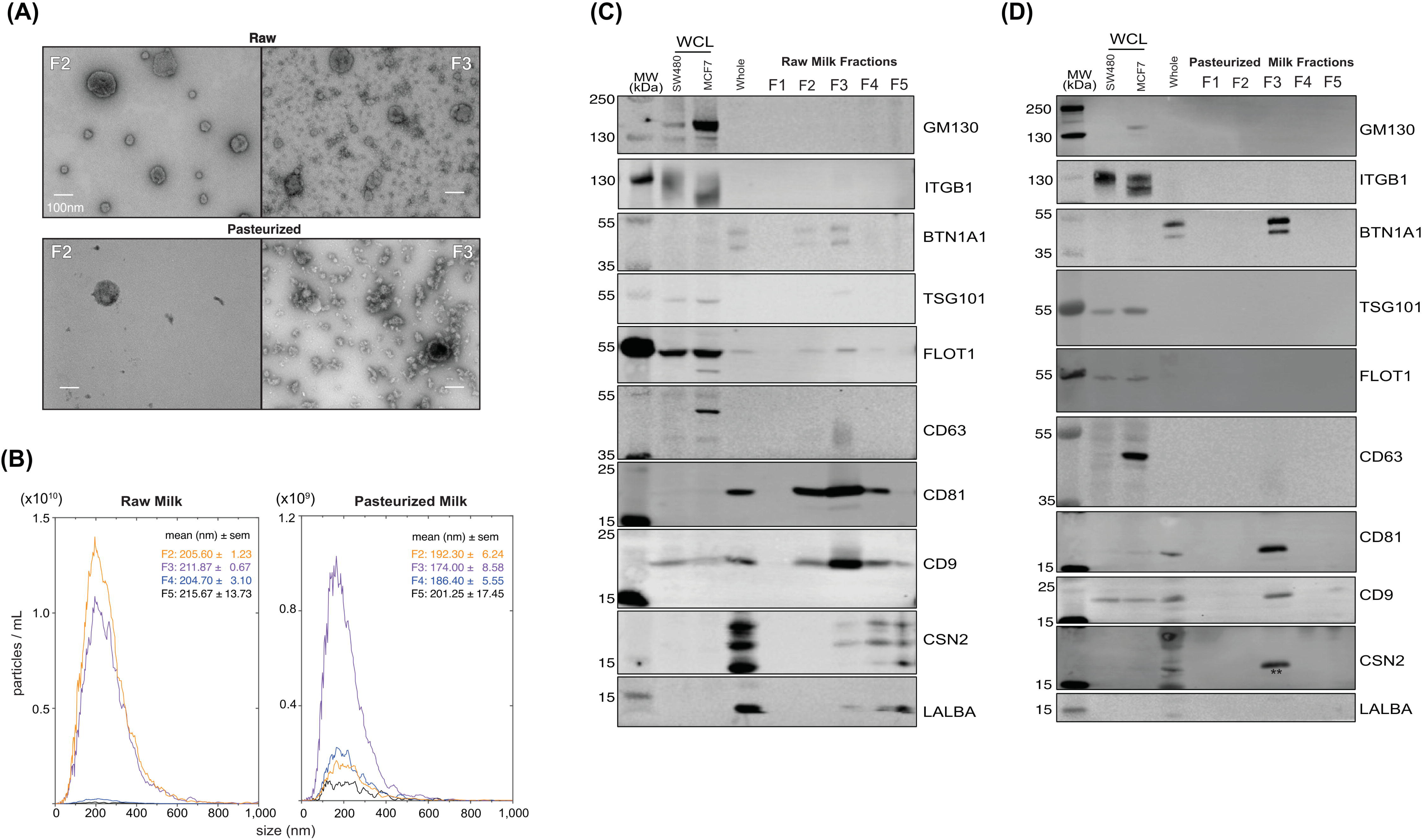
Pipeline effectively isolates human milk extracellular vesicles (HMEVs) from raw (RHM) and pasteurized donor human milk (PDHM). All data shown are from the same RHM or PDHM sample isolated using the pipeline in Figure 1. (A) Transmission electron micrographs of fractions (F) 2 and 3 from RHM and PDHM. Images represent data collected across n=3 biological replicates obtained from three different donors and three different donor human milk sources (**Table 1**). Scale bar = 100 nm. (B) Nanoparticle tracking analysis (NTA) was performed on RHM and PDHM F2-5. F1 was below the level of detection. Data represent the average of triplicate runs for each fraction and data is smoothed using GraphPad Prism® v10. Size data represents the average median size across triplicate runs. (C) Western blotting assessed the protein profile of isolated HMEVs. Fifteen micrograms of SW480 (colorectal cancer cells), MCF7 (breast cancer cells) whole cell lysates (WCL), and whole milk were loaded as positive controls. Twenty microliters of unconcentrated fractions (F) 1-5 of RHM (C) and PDHM (D) were run on a 4-12% Bis-Tris gel, transferred to a PVDF membrane, and probed according to MISEV 2018 guidelines, including controls for common proteins found in human milk (BTN1A1, LALBA, CSN2). Each blot represents triplicate experiments performed on three biological replicates of RHM and PDHM. ** indicates residual CD81 signal from re-probing the membrane for CSN2 and using the same color secondary.

Western blotting of EV proteins followed the MISEV2018 guidelines (13) to confirm EV enrichment and purity based on the presence and absence of indicated marker proteins. F2 and F3 isolated from RHM (**Figure 2C**) and F3 isolated from PDHM (**Figure 2D)** were enriched for transmembrane proteins CD9, CD63, and CD81. Cytosolic proteins TSG101 and FLOT1 were recovered within RHM EVs, but absent from the pasteurized samples. Our samples lack microvesicle protein ITGB1 (49). Importantly, all of our samples lack the Golgi protein GM130. The RHM sample is depleted of the whey protein LALBA and highly abundant milk protein CSN2 (**Figure 2C**), while PDHM sample lacks LALBA and CSN2 (**Figure 2D**). Note, the CSN2 band marked by ** in **Figure 2D** is CD81 signal as the blot was re-probed using the same color secondary. True CSN2 signal is a multi-band pattern as shown in the whole milk positive control. Additionally, the RHM and PDHM samples also contain BTN1A1, a protein that is prevalent in milk fat globules (56, 57), but also potentially present in HMEVs, as demonstrated by proteomics (58) (**Figure 2C & D**). The western blots support the TEM and NTA data showing EV markers in F2-3 of RHM and only F3 of PDHM. Notably, these data suggest that pasteurization may deplete cytosolic proteins, such as TSG101 and FLOT1.

Collectively, these data demonstrate that our pipeline effectively and consistently isolates EVs from small volumes of RHM and PDHM.

### 3.3 Di8-ANEPPS-positive particles are disrupted by Triton-X

Next, we wanted to validate the specificity of our Di8-ANEPPS staining and the level of non-membranous particles in our sample preparations. First, we determined the total number and size of all particles non-fluorescently using scatter detection mode This includes membrane and non-membrane bound particles, as well as microbubbles. The Zetaview NTA uses scatter for particle detection only; the sizing is measured by tracking the Brownian motion of detected particle and applying the Stoke Einstein’s equation (59). Notably, the total number of particles detected by scatter was quite high relative to the Di8 signal, suggesting that Di8 excludes a large proportion of background particles, potentially due to the exclusion of nonmembrane-bound particles. To further examine the ability of Di8 to detect non-membranous particle contamination (protein aggregates or casein micelles), we tested the sensitivity of F2 and F3 to Triton-X100, EDTA, and proteinase K (**Figure 3**). We found that of particles labeled with Di8 in F2 and F3 the majority (>50%) are Triton-X solublizable, consistent with EVs (60), as seen by the reduction in particle counts with the addition of Triton-X. Importantly, each fraction contained some particles that were sensitive to EDTA or proteinase K, which were detected by Di8 and therefore also counted in our particle measurements. This means that when using Di8 to quantify HMEVs by NTA, the actual number detected can be augmented slightly by non-membranous particles. Importantly, the use of Di8 excludes the vast majority of non-membrane bound or non-specific particles detected by scatter alone. Based on this experiment, F3 seems to contain membranous/EV particles versus F2.

**Figure 3.**
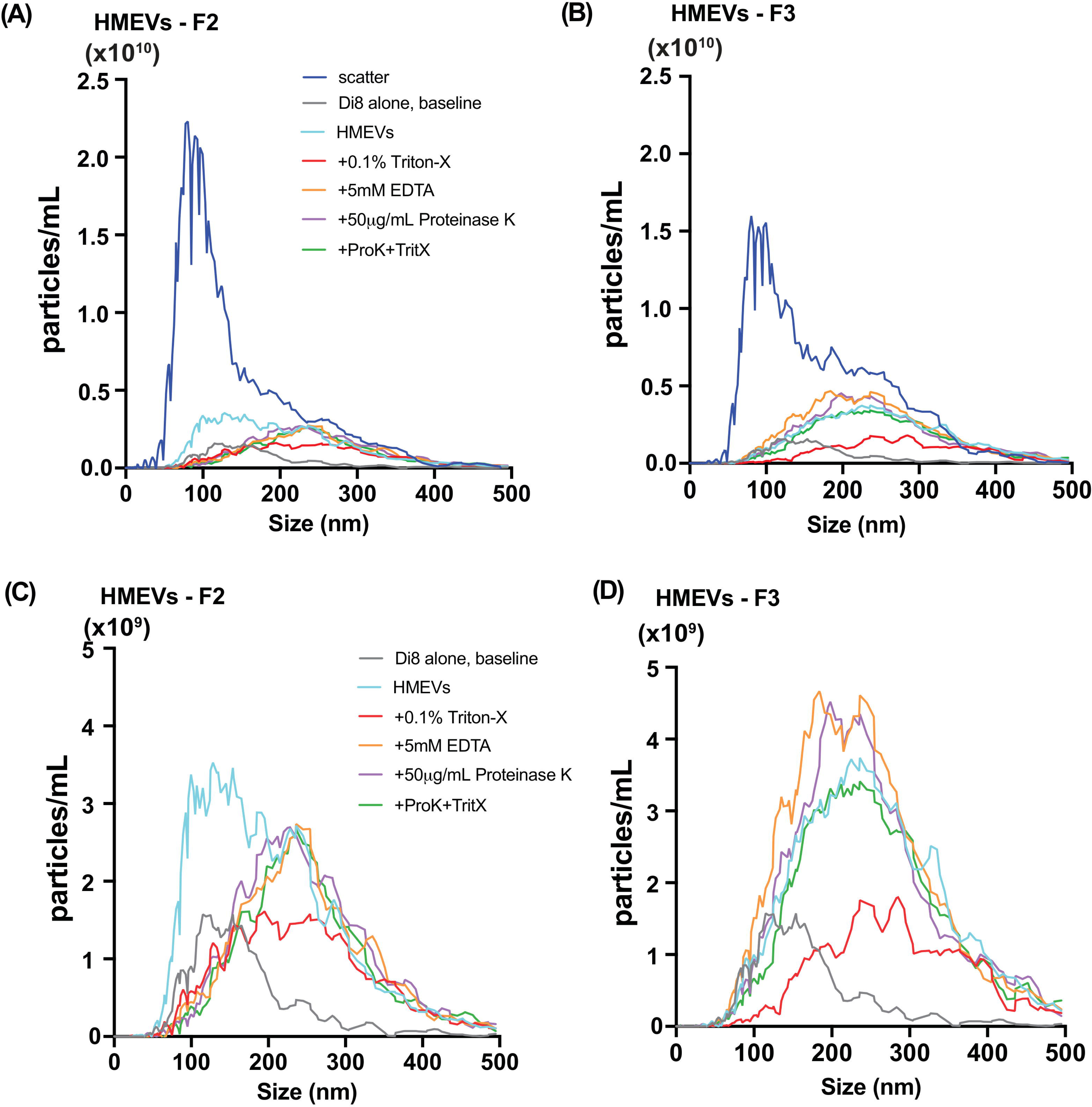
Di8 ANEPPS detects the majority of Triton-X-solublizable EVs in fraction 3 from human milk. Fractions 2 (A, C) and 3 (B, D) were isolated from human milk sample (c), concentrated using Amicon^®^ concentration column (10kDa MWCO), and labeled with Di8-ANEPPS. Aliquots of the labeled sample were then incubated with either 0.1% Triton-X100 (final concentration), 5mM EDTA (final concentration), 50 µg/mL proteinase K (Thermo Fisher Scientific, final concentration), or 0.1% Triton-X100 + 50 µg/mL proteinase K before particle counts were measured by nanoparticle tracking analysis (NTA). (A & B) show the Di8 particle measurements for comparison with scatter, while (C & D) show measurements detected with the fluorescence reading alone. Data represent the average of triplicate runs for fractions 2 and 3 of a single milk sample. Data are smoothed using GraphPad Prism® v10.

### 3.4 EVs numbers decrease within the neonatal small intestine

Neonatal human digesta was collected by gravity after gastric feeding from oro- or nasoduodenal/jejunal sampling tubes or ostomy bags from three neonates in the OHSU NICU (**Figure 4A**; **Table 2**). Infants were fed RHM and digesta was isolated from the proximal or mid-small intestine (**Figure 4B)**.

**Figure 4.**
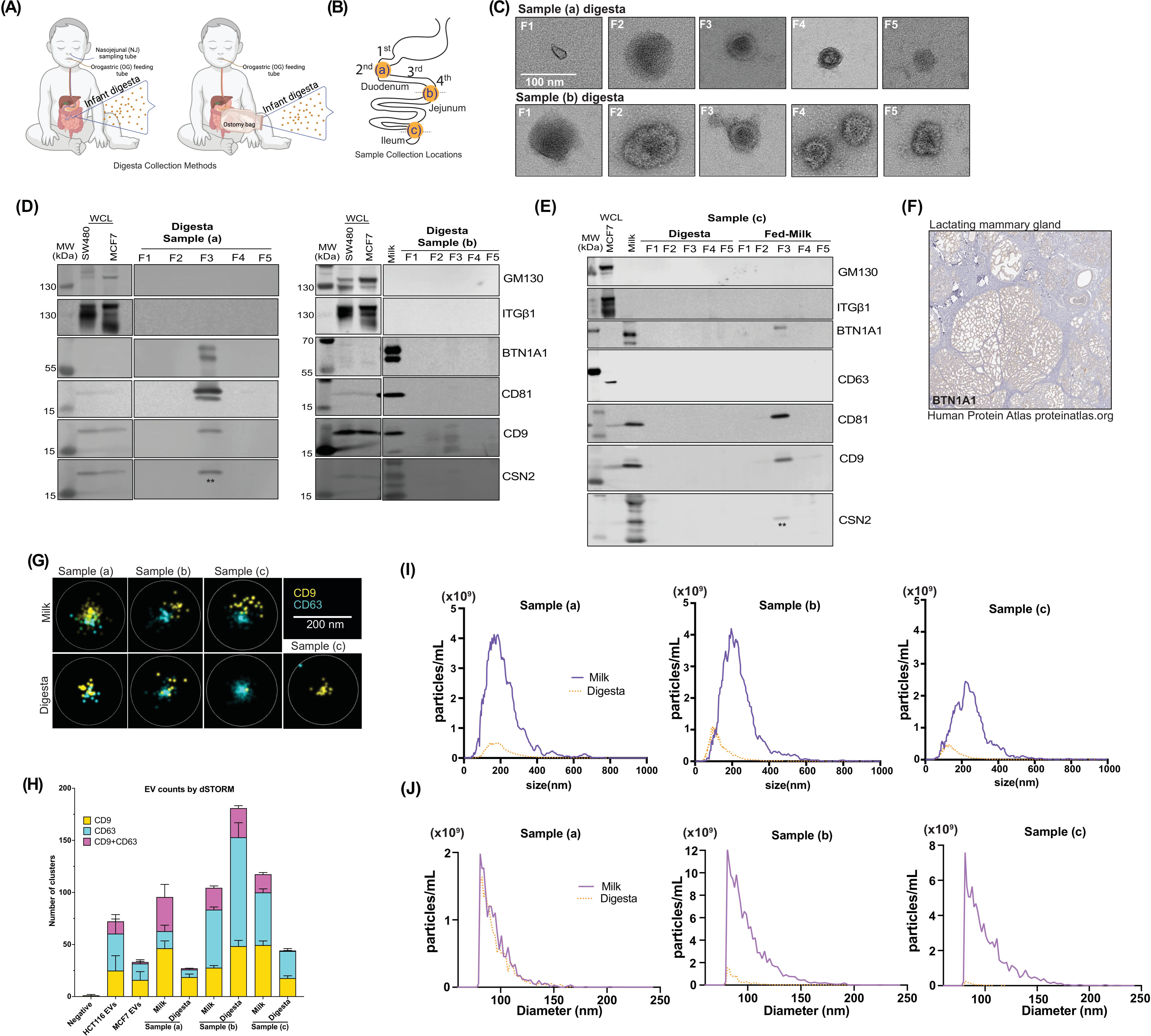
Pipeline effectively isolates extracellular vesicles from neonatal human digesta (dEVs). (A) Neonatal human digesta was collected from infants in the NICU after obtaining parental informed consent. Inclusion criteria included infants already admitted to the NICU, greater than 26 weeks corrected gestational age with an indwelling nasogastric or orogastric feeding tube who were tolerating full enteral feeding volumes. Infants were excluded from the study if they had anatomic or functional gastrointestinal disorders, were medically unstable, were non-viable, or had disorders that would be expected to affect normal digestion. Before feeding, a nasal tube was placed into the proximal small intestine. Feeds were delivered via a gastric tube over 30 min or less. (B) Data are from three participants with sampling tubes placed along the duodenum (a, b) or from an ostomy (c), as shown. (C) Transmission electron micrographs (TEM) of fractions (F)1 – 5 of neonatal digesta collected from participants (a) and (b). Scale bar = 100 nm. (D) Western blotting assessed the protein profile of isolated dEVs in the participants (a), and (b) or (E) the digesta and milk samples from participant (c). Fifteen micrograms of SW480 (colorectal cancer cells), MCF7 (breast cancer cells) whole cell lysates (WCL) and/or whole milk were loaded as positive controls. Each digesta fraction was concentrated over an Amicon^®^ concentration column (100kDa MWCO) to allow for loading of the entire fraction volume in the gel; therefore, the data shown represent all the proteins in the fraction from a 1 mL digesta sample. Samples were loaded on a 4-12% Bis-Tris gel, transferred to a PVDF membrane, and probed according to MISEV 2018 guidelines, including controls for common proteins found in human milk (BTN1A1, LALBA, CSN2). LALBA and CSN2 were not detected. To preserve the samples, membranes were re-probed for some proteins. ** indicates residual CD9 signal from re-probing the membrane for CSN2. CSN2 was not detected in EV samples, see banding pattern in whole milk positive control in participants’ (b) and (c) blots for comparison. ITGβ1 was detected by re-probing the membrane after GM130. Both antibodies are rabbit, resulting in the smear in control WCL lanes for ITGβ1. The isolated fractions were all negative. (F) Immunostaining for BTN1A1 in lactating mammary gland. Image provided courtesy of Human Protein Atlas (54, 55). (G) Representative images of direct stochastic optical reconstruction microscopy (dSTORM). Human milk and digesta EV samples were affinity captured on microfluidic chips using CD9/CD81/CD63 antibodies and labeled with CD9-CF488 (excitation (ex)/emission (em): 490/515 nm) and CD63-CF568 (ex/ em: 562/583 nm) antibodies for imaging on the Nanoimager S Mark II microscope (ONI) with 100X oil-immersion objective and freshly prepared dSTORM-imaging buffer. Data were processed on NimOS software (version 1.19, ONI). Subpopulation analyses of EVs that express one or two markers were analyzed using ONI’s online platform CODI (https://alto.codi.bio). We used a density-based clustering analysis with drift correction and filtering to evaluate each vesicle. At least 5 localizations were required to constitute a vesicle and at least 3 localizations of one protein were required to consider the localizations a real signal. Data were compiled in Microsoft Excel™ and (H) the localizations within each sample were summarized using GraphPad Prism® v10. (I) Nanoparticle tracking analysis (NTA) was performed on matched fed-human milk and neonatal digesta F3 isolated from participants (a), (b), and (c) and stained with Di8-ANEPPS. Data represent the average of triplicate runs for each sample and the data is smoothed using GraphPad Prism® v10. (J) Resistive pulse sensing measurements were performed on matched fed-human milk and neonatal digesta F3 isolated from participants (a), (b), and (c). EV samples were diluted 1:20 in 1% Tween-20 (v/v) in 0.2µm filtered PBS and loaded in polydimethylsiloxane cartridges (diameter range 65 nm to 400 nm, C-400). From each 4 μL sample, four thousand events were collected using a Spectradyne nCS1™ instrument and analyzed using the accompanying nCS1™ Data Viewer (Version 2.5.0.325). Size and concentration measurements were calibrated based on 150 nm beads. Concentrations were adjusted using Microsoft Excel™ and graphs were made in GraphPad Prism^®^ v10.

We isolated and validated the dEVs identically to the HMEVs described above. Negative TEM shows the greatest consistent, enrichment of cup-shaped EV structures in F3 (**Figure 4C, Supplemental Figure 1**). This is further supported by western blotting showing enrichment of EV markers CD9 and/or CD81 in F3 of digesta samples (a) and (b) (**Figure 4D**). No EV markers were detected by western blotting in digesta sample (c), although they are present in the fed-milk sample (**Figure 4E**). Western blotting followed the MISEV2018 guidelines (13) to confirm EV enrichment and purity based on the presence or absence of indicated marker proteins. All the samples lacked ITGB1, GM130, and CSN2 (**Figures 4D & 4E**). The bands in the CSN2 blots marked by ** are residual CD9 signals as the blots were re-probed for CSN2. We detected BTN1A1 prominently in digesta sample (a), faintly in digesta sample (b), and in fed milk from sample (c), suggesting that BTN1A1 may be present within some dEVs or that isolates contain milk fat globule membrane components. Notably, BTN1A1 is only expressed in the lactating mammary gland (**Figure 4F**) and nowhere else along the digestive tract, including the salivary gland, pancreas, gallbladder, or liver (55), and was previously found within HMEVs by mass spectrometry (58).

We further confirmed the EV identity of our samples using super-resolution microscopy (**Figure 4G**). Particles were captured with CD9/CD81/CD63 and subsequently labeled with CD9 and CD63 antibodies. We detected single-labeled EVs in all samples and dual-labeled EVs in all but sample (c) digesta (**Figures 4G and 4H**).

Based on these data, the minimal impact of EDTA or proteinase K on F3 EVs (**Figure 3**), and the higher protein levels in F3 vs F2 (**Supplemental Table 1**), we focused on F3 as the EV-containing fraction in subsequent analyses (**Table 3**), NTA using Di-8-ANEPPS lipid membrane dye (**Figure 4I**), and resistive pulse sensing (RPS) (**Figure 4J**). The dEV protein concentrations were ∼2-fold lower and particle counts by NTA were ∼3-8-fold lower than the isolated HMEVs from the input milk samples (**Table 3**; **Figure 4I**). The dEV particle counts measured by RPS were lower for samples (b) and (c), but not for sample (a) (**Figure 4J**).

**Table 3.** Matched milk and digesta EV protein concentrations.

|  | Protein (ng/uL) |  |  |
| --- | --- | --- | --- |
|  | sample (a) | sample (b) | sample (c) |
| Intestinal Region | Proximal | Proximal | Middle |
| **Fed Milk** | 82.83 | 215.77 | 86.47 |
| **Digesta** | 41.64 | 100.79 | 45.30 |
| **Fold-change** | 1.99 | 2.14 | 1.91 |

Note: Data is also shown in Supplemental Table 1 for comparison with F2 data.

Collectively, these data indicate that dEVs can be isolated from neonatal digestive fluid and that dEV amounts are generally less than the input milk. Since there is no validated marker unique to HMEVs, we cannot confirm that these isolated EVs are from human milk alone, therefore we term them digesta EVs (dEVs).

### 3.5 Neonatal human enteroids take up dEVs

We established that dEVs can be isolated from infant digesta. Next, we asked whether neonatal IECs take up dEVs. To test this, we developed a neonatal human enteroid model. Patient-derived enteroids, enteroid-derived monolayers, or gut-on-a-chip systems are emerging models for studying neonatal intestinal physiology and pathophysiology (61–65). Our model is derived from intestinal epithelial stem cells isolated from a 2-month-old participant undergoing surgery for ileal atresia.

Our neonatal enteroids were propagated in the basal-out orientation where they grow in Matrigel and the basolateral cell surface faces the culture media (**Figure 5A**). For these studies, we used the apical-out enteroid model where the enteroids are cultured in suspension in the absence of a basement membrane (52, 61, 64). This leads to polarity reversal, shown by phalloidin staining (green) on the outer surface of the enteroid (**Figure 5A**). The apical-out orientation allows for direct access to the apical cell surface, making the culture media effectively equivalent to the intestinal lumen and allowing for manipulation of luminal exposures, such as dEVs. We first analyzed gene expression for IEC-type specific markers, demonstrating that our apical-out enteroids significantly downregulate markers of proliferation *Ki67* and *LGR5* and upregulate more differentiated markers, such as *CHGA* relative to basal-out enteroids (**Figure 5B**), which are generally more proliferative (52). Notably, our neonatal enteroids lack Paneth cells (*DEF6*) at baseline, consistent with published findings (66), and down-regulate *BEST4* mRNA. This suggests that our apical-out enteroids represent a more differentiated and less proliferative intestinal epithelium.

**Figure 5.**
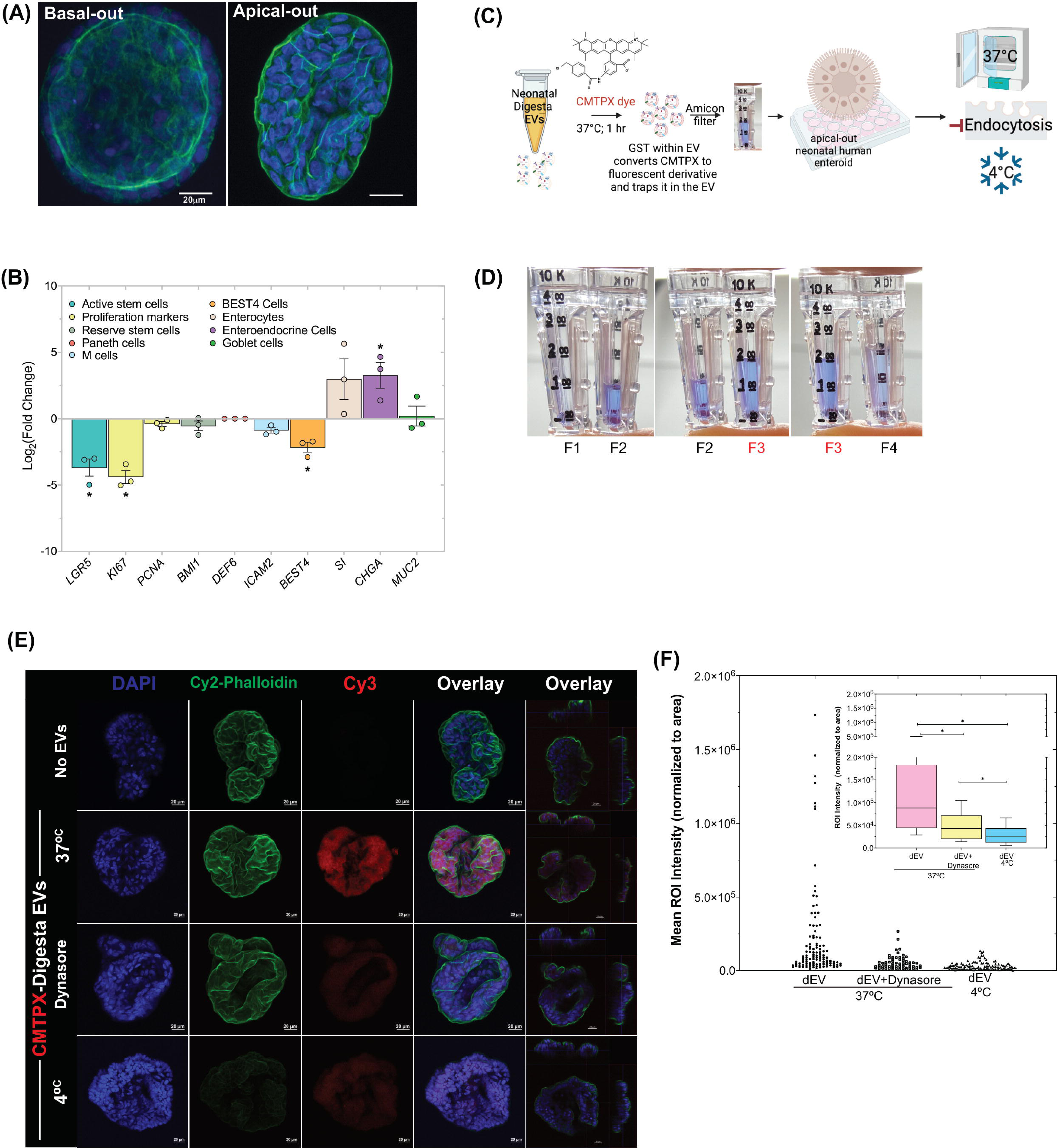
Neonatal human enteroids take up neonatal digesta extracellular vesicles (dEVs) primarily by dynamin-mediated endocytosis. (A) Neonatal human enteroids were cultured in Matrigel basement membrane where the basal cell surface faces the media (basal-out) before removing the basement membrane to induce polarity reversal to the “apical-out” orientation for all experiments. DAPI (blue) stains nuclei and phalloidin (green) stains F-actin. Scale bar = 20 µm. (B) Polarity reversal and the apical-out orientation down-regulated expression of *LGR5* and *KI67* relative to basal-out controls. *BEST4* was down-regulated, whereas *CHGA* was increased across passages. Each dot represents a separate passage (biological replicate). Gene expression values are the average of triplicates for each gene within each biological replicate and are normalized to the housekeeping gene *RPLP0*. Data are expressed at Log2-fold change and error bars represent the standard error of the mean (SEM). \**p*<0.05 was considered statistically significant. (C) dEVs were isolated from samples (a) and (b) and stained with 10 µM CMTPX dye in a 37 °C water bath for 1 hour in the dark. (D) The excess dye was removed using Amicon^®^ Ultra-0.5 10 kDa MWCO centrifugal filters and samples were concentrated to the desired volume for uptake experiments. Apical-out enteroids were treated with CMTPX-stained EVs for 90 minutes at 37 °C in the presence or absence of the endocytosis inhibitor Dynasore or at 4 °C. The “No EV” control consisted of PBS incubated with CMTPX dye and filtered through an Amicon^®^ filter. (E) EV uptake was assessed by confocal microscopy on fixed whole-mount enteroids. DAPI (blue) marks nuclei. Phalloidin (Cy2-green) marks F-actin. CMTPX staining is shown in Cy3-red. Scale bar = 20 µm. Data represent a single experiment using a pool of dEVs from participants (a) and (b). Images are representative of enteroids imaged across 4 slides for each treatment group. Images were acquired and processed using identical microscope settings. Notably, the Cy2 intensity for the phalloidin staining in the xz.yz projection image for the 4°C treatment was increased 3-fold to enable visualization of the cell surface. All other channel intensities are matched. (F) CMTPX-labeled dEV uptake was quantified using images captured with a 2X objective on a Keyence BZ-X800 microscope. Staining intensity within each enteroid was determined using ImageJ and normalized to the enteroid area. Data represent individual enteroids across 4 slides in each treatment group from a single experiment. The inset shows box and whisker plots of the same data. Data was analyzed by one-way ANOVA with Dunn’s test for multiple comparisons. \**p*<0.05 was considered statistically significant.

Once established and validated, we asked whether neonatal enteroids readily take up dEVs. Since our dEVs are isolated from participant samples, we are unable to label and track using fusion proteins, or fluorescent tags (32, 67). Instead, we developed a novel method for labeling the dEVs using CMTPX dye (**Figure 5C**). This dye is nonfluorescent initially but readily permeates the EV lipid bilayer where it is then converted to a fluorescent derivative by glutathione-S transferase (GST) and retained within the EV. This is important for demonstrating the specificity of EV uptake. We observed the highest level of CMTPX staining in fraction 3 (F3) consistent with TEM, NTA, and western blot data showing the highest number of HMEVs or dEVs in F3 (**Figures 2, 3, and 4**).

All images were acquired using identical microscope settings. Representative images are shown in (**Figure 5E**). dEV uptake was quantified using ImageJ (**Figure 5F**). At 37 °C we observe robust uptake of CMTPX-labeled dEVs by neonatal human enteroids as indicated by a strong red signal. The specificity of the labeling and the uptake were confirmed using a dynamin-2 inhibitor (Dynasore) to block clathrin and caveolin-dependent endocytosis and by performing experiments at 4 °C to block all forms of energy-dependent endocytosis (32, 67–70). Collectively, our results indicate that neonatal human enteroids rapidly take up dEVs primarily via energy-mediated endocytic processes, while a smaller proportion of dEVs are taken up in the presence of dynamin-2 inhibition. Incubation at 4 °C inhibits nearly all uptake. Further, the data indicate that our labeling is specific and that our samples contain low to negligible levels of unincorporated CMTPX, as the free dye would show up as labeling via diffusion in the Dynasore + 37 °C or 4 °C samples.

## 4. DISCUSSION

HMEVs are emerging therapeutics and drug delivery vehicles (71–78). Cell culture and animal model studies indicate that undigested, unmodified HMEVs may affect intestinal epithelial biology (15, 19, 23), and limit damage from necrotizing enterocolitis (16, 17, 20–22). Further studies indicate that HMEVs survive in vitro simulated digestion (15, 24, 25), may transit through an animal’s digestive tract to transfer cargo or affect other organs, including the brain (15, 28), and/or enhance cognitive performance (29). However, whether these nano-sized information carriers survive digestion to reach the human intestine remained an open question. To address this knowledge gap, we obtained in vivo digested milk samples and optimized an isolation protocol capable of isolating HMEVs from small sample volumes.

Isolating EVs from human milk is challenging due to protein complexes, such as casein micelles, immunoglobulins, and lipoproteins that can entrap HMEVs, mask less abundant proteins, and interfere with downstream analyses, such as western blotting or mass spectrometry (79). Protein complexes can also mimic the appearance and size of EVs in NTA or RPS. To circumvent these pitfalls, it is common to perform HMEV isolations on large starting volumes, which allow for multiple processing steps to enrich for EVs and remove other bioactive factors that may complicate data interpretation (15, 17, 18, 46, 80, 81). This can be prohibitive if sample volumes are limited (≤ 2mL). In such cases, some published studies use precipitation-based methods, such as ExoQuick^®^ and ExoEasy^®^ (82–84). Although these methods are efficient, they can result in co-isolation of non-EV proteins (85–88). Additionally, EV fractions obtained from ExoQuick^®^ may contain traces of biopolymers that can interfere with mass spectrometry (89).

Herein we optimized an isolation pipeline developed for large volumes of milk (46) for use with a 1 mL starting volume of human milk or digesta. This method utilizes a combination of acetic acid precipitation and density gradient ultracentrifugation to achieve relatively high yield and relatively high purity HMEVs. We validated our methodology using RHM and PDHM. Our RHM samples were consistently enriched for a combination of tetraspanins CD9, CD63, and CD81 and intravesicular markers FLOT1 and TSG101 (13) and de-enriched for microvesicle marker ITGB1 (49), Golgi marker GM130 (13), and milk-specific proteins CSN2 and LALBA (90). CD63 shows up as heterogeneous in size in our HMEV samples. This is potentially due to the heavy glycosylation of milk proteins (91). Notably, samples were also enriched for BTN1A1. BTN1A1 is exclusively expressed in lactating mammary glands (55) and prevalent in milk fat globules (56, 57). It was previously detected in HMEVs by mass spectroscopy (58), suggesting its potential as a HMEV marker. To demonstrate that BTN1A1 is a marker of HMEVs super-resolution microscopy or immunogold-labeled TEM is required.

To further validate our isolation pipeline, we compared raw HMEVs with those isolated from PDHM. Pasteurization most commonly heats milk to 63° C for 30 minutes to reduce bacterial load (pathogenic and nonpathogenic) and is currently the standard method of human milk processing to ensure safety (92). Pasteurization, however, damages, destroys, or alters many bioactive factors found in milk (93). This is consistent with our findings. We observed that pasteurization resulted in a depletion of tetraspanin proteins and cytoplasmic EV protein markers (FLOT1, TSG101). This is consistent with the published work of others showing that pasteurization reduces CD63 protein within human (94) and bovine milk EVs (95). However, there may be some variation between species or with processing methods as one study found TSG101 in pasteurized bovine milk (96). Notably, our raw and pasteurized samples are not matched from the same parental source, limiting our ability to draw firm conclusions about the effects of pasteurization on EV size, number, or cargo. Previously published reports indicate that size and number are unchanged by pasteurization (94). Our data add to a growing body of evidence demonstrating that pasteurization could damage or destroy some populations of HMEVs (94, 95, 97). Additional research is needed to fully understand the effects of milk processing on HMEVs.

Our isolation pipeline also effectively isolated EVs from human milk-fed neonatal digesta (dEVs), enabling us to demonstrate that EVs are present in the intestine of human milk-fed neonates. These EVs could be derived from the fed human milk, secreted by cells within the digestive tract, or a combination of the two. In vitro-simulated digestion studies suggest that HMEVs can survive low pH and the presence of some digestive enzymes (15, 24, 25), supporting the hypothesis that some of our dEVs may be from human milk. To define the precise source of the dEVs we need specific markers that are exclusively present within the lactating mammary gland, such as BTN1A1, or the neonatal intestinal epithelium. This study does not directly define BTN1A1 as an HMEV marker; therefore, we cannot determine the precise origin of our dEVs. Nonetheless, our data indicate that a lactating mammary gland-specific protein is present in some dEV samples isolated from milk-fed infants meaning that either HMEVs or other human milk components are co-isolating with the dEVs from these participants. The absence of BTN1A1 in digesta sample (c) may be due to the distal sampling location (jejunostomy) relative to samples (a) and (b). Across the three samples examined, the level of dEV protein decreased by a factor of ∼2 between fed human milk and the intestinal digesta. The number of dEVs decreased by 3-8-fold relative to HMEVs as measured by NTA. RPS measurements show equivalent or greater decreases in particle number for samples (b) and (c), while sample (a) measured similarly to the input milk sample. This could be due to non-membranous particles in the digesta inflating the measurement or a falsely low RPS measurement for the fed milk sample. Taken together, across protein concentration and multiple methods of particle measurement, this suggests that dEVs are likely reduced within the intestinal digesta compared to the fed milk sample.

The reduction in dEVs relative to HMEVs could be due to several factors: 1) sample dilution with digestive secretions, 2) EV destruction during transit and digestion, 3) cellular uptake, or 4) a combination. Previous work from the Dallas lab indicates that intestinal secretions result in 1.15-1.5-fold dilution of digesta in the intestine relative to the input milk as determined by measuring the concentration of indigestible PEG-28 in fed-human milk and infant digesta using mass spectrometry (98). This suggests that the observed decrease in dEVs could be partly due to dilution from digestive fluids. However, the reduction that we observe in protein concentration or particle number is greater than what would be expected with dilution alone (**Table 3**), indicating that HMEVs are either rapidly destroyed or absorbed by the digestive tract. Our uptake data indicate relatively rapid dEV uptake, suggesting that uptake may also contribute to reduced dEV numbers within the neonatal small intestine. Notably, if neonatal intestinal EVs are present in our samples, these would only augment the signal and further suggest that HMEVs are reduced within the digesta. Definitive EV biomarkers are needed to determine the origin(s) of the dEVs. Further research is also needed to fully address differences between HMEV and dEV numbers, including normalization to an indigestible substance such as PEG-28 to account for the effects of dilution.

To better understand the specificity of the Di8 labeling in our human milk samples, we tested the effects of Triton-X, EDTA, and proteinase K on our NTA measurements. These results reveal that although Di8-labeling is an imperfect method of measuring HMEVs, it is far superior than scatter alone, which detects a much higher background of non-specific signal. Although the majority of the particles measured were Triton-X solublizable, the quantification was altered by the addition of EDTA or proteinase K to the samples. As a result, the NTA measurements from our samples are not exact counts of EVs, but rather an approximation of the particle concentration including HMEVs and some non-membranous particles. The number of non-membranous particles is most readily appreciated when comparing the NTA data in (**Figure 4I**) with the RPS data (**Figure 4J**), as well as in **Figure 3** comparing scatter versus Di8 labeling in F2 and F3. F2 especially contains many non-membranous particles, some of which are still detected by Di8 and altered with Triton-X, EDTA, or proteinase K. Each method of EV quantification has limitations (99), however, having performed several different methods, protein concentration and particle measurements via NTA and RPS, the collective data suggest that dEVs are generally reduced in number relative to the HMEVs found within the matched milk sample.

Once we isolated dEVs, we next asked whether they were taken up by non-transformed, primary neonatal IECs. To do this, we used a neonatal enteroid (intestinal organoid) model for dEV uptake. These enteroids grow rapidly and express expected markers of proliferation in the basal-out state. We utilized published protocols (52, 64) to induce polarity reversal in the enteroids allowing direct access to the apical surface from the culture media. We confirmed the apical-out orientation with phalloidin-staining and gene expression showing a significant down-regulation in proliferative markers, consistent with published studies (52). Notably, markers of absorptive (*SI*) and secretory lineages (*CHGA* and *MUC2*) were elevated, but only differences in *CHGA* were statistically significant, suggesting some variability in the cell lineages present within the apical-out enteroids.

We then used CMTPX dye to track dEVs ex vivo. Nonfluorescent CMTPX freely diffuses into HMEV double-layered membranes. Once inside, it undergoes a chemical transformation mediated by GST (100) rendering it fluorescent and trapping it within the EV. This allows for extra-EV CMTPX to be washed away using chromatography column treatment. Therefore, CMTPX specifically labels isolated EVs and avoids non-specific labeling of cellular or organelle membranes often seen with lipophilic dyes, such as PKH26 (red), PKH27 (green), and C5-maleimide-Alexa633 (100, 101). We observed minimal labeling when endocytosis was inhibited chemically and when all uptake mechanisms were limited by cold temperatures, suggesting that our labeling method is specific and indicates minimal signal from passive dye diffusion. This indicates that CMTPX may be a superior non-genetic method for labeling and tracking EVs.

Our data suggest that dEVs are rapidly taken up by neonatal enteroids in a process that depends in part on dynamin-mediated endocytosis. This is consistent with the work of others demonstrating that bovine milk EVs are transported via endocytosis into endothelial cells (68) or transformed Caco2 or IEC6 cells (70). In our study, uptake was further significantly decreased by incubation at 4°C, suggesting that other mechanisms may also be involved in dEV uptake within the intestine. These could include lipid raft/caveolae-mediated endocytosis (102), phagocytosis, and/or micropinocytosis (103). Importantly, our studies establish a relevant system for performing future functional studies and build on published work in immortalized cell lines (24, 25) to show that non-transformed neonatal enteroids readily absorb dEVs. Additional studies are needed to demonstrate that dEV uptake results in functional changes in enteroid physiology or gene expression.

Collectively, this study demonstrates that dEVs can be isolated from infant intestinal contents post-human milk feeding. These dEVs can then be rapidly absorbed ex vivo by neonatal IECs in a process that is partially dependent on dynamin-mediated endocytosis. This study provides a modified protocol for use with small sample volumes and lays the foundation for future, in-depth work investigating biomarkers of human milk and digesta EVs, the effects of donor human milk processing and gastrointestinal digestion on HMEV structure and contents, and how dEVs interact with and are taken up by the neonatal intestinal epithelium.

We chose the more inclusive term extracellular vesicles (EVs) for the particles isolated in this study, as we did not directly examine their biogenesis. However, our data suggest that they are most consistent with exosomes or smallEVs based on our density-gradient ultracentrifugation method of isolation, the particle size, and the presence of tetraspanin proteins CD63, CD81, CD9, and ESCRT protein TSG101 (13, 104).

## FIGURE LEGENDS

**Supplemental Figure 1.**
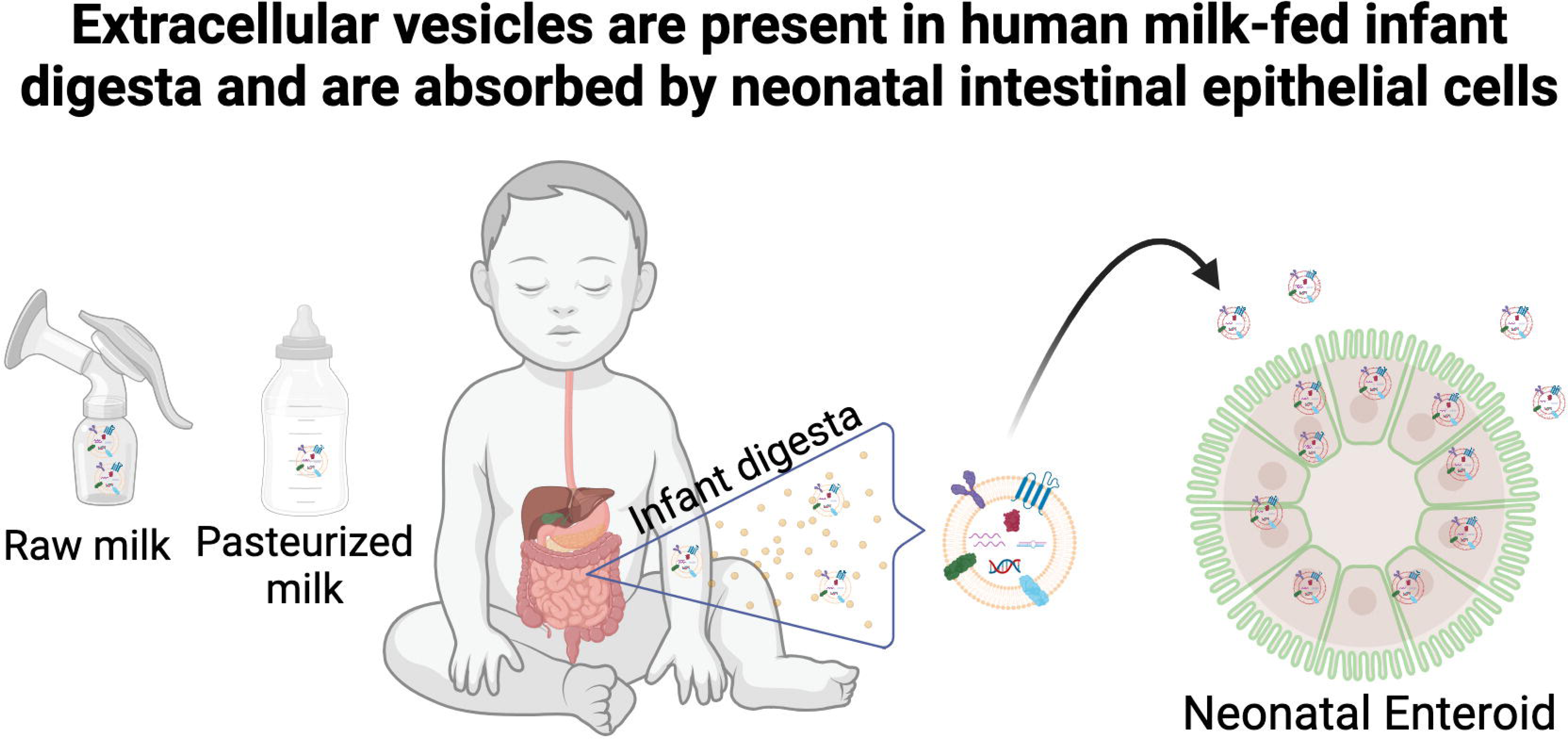
Transmission electron micrographs (TEM) of fractions 1-5 isolated from human milk-fed infant digesta. Transmission electron micrographs of fractions (F)1 – 5 of neonatal digesta collected after human milk feeding from participants (a) and (b). Scale bar = 100 nm. Higher magnification TEM images from each fraction are shown in Figure 4C.

## AUTHOR CONTRIBUTIONS

**Conceptualization**: Sarah F. Andres, Brian Scottoline, Randall Armstrong

**Sample acquisition and participant data management:** Brian Scottoline, Amy Olyaei Molly Aloia

**Methodology**: Yang, Zhang, Madeline Kuhn, Randall J. Armstrong, Sarah F. Andres

**Data acquisition:** Claire Yung, Yang Zhang, Madeline Kuhn, Sarah F. Andres, Randall Armstrong

**Funding acquisition**: Sarah F. Andres, Brian Scottoline

**Writing**: Sarah F. Andres, Yang, Zhang, Claire Yung, Madeline Kuhn, Brian Scottoline

**Review & editing**: Sarah F. Andres, Brian Scottoline, Randall Armstrong

**Approved manuscript:** all authors

## ACKNOWLEDGEMENTS

SFA is supported by the National Institutes of Health (NIH) grant K01DK129401 and grants from the Collins Medical Trust, Medical Research Foundation of Oregon, and OHSU Exploratory Research Seed Grant. BS is supported by the Gerber Foundation #21-9161, a grant from Evolve BioSystems, and NIH grant R01HD097367. The authors would like to thank Drs. Pepper Schedin and Reuben Hoffmann for sharing the MCF7 cell line. The authors kindly acknowledge expert technical assistance from: Steven Adamou, Erin Stempinski, and Dr. Claudia Lopez from the OHSU Multiscale Microscopy Core for their technical expertise and assistance with the negative staining and electron microscopy; Brian Jenkins, Dr. Felice Kelly, and Dr. Stefanie Kaech Petrie from the OHSU Advanced Light Microscopy Shared Resource for their expert technical expertise and assistance with confocal microscopy; Dr. Julie Saugstad, Trevor McFarland, and Sarah Catherine Baker for their help with the resistive pulse sensing data acquisition and analysis; Vanessa Lambatan for help with super-resolution imaging and analysis.

## DATA AVAILABILITY STATEMENT

The data that support the findings of this study are available from the corresponding author upon reasonable request.

---

**Supplemental Table 1.** Matched milk and digesta F2 and F3 protein concentrations.

|  | Protein (ng/uL) |  |  |  |  |  |
| --- | --- | --- | --- | --- | --- | --- |
|  | sample (a) |  | sample (b) |  | sample (c) |  |
|  | F2 | F3 | F2 | F3 | F2 | F3 |
| **Fed Milk** | 11.89 | 82.83 | 32.55 | 215.77 | 20.44 | 86.47 |
| **Digesta** | 6.76 | 41.64 | 69.60 | 100.79 | 16.42 | 45.30 |

Note: F3 data is also shown in Table 3 and provided here for comparison.

