## Supplementary figures and images for "Neonatal enteroids absorb extracellular vesicles from human milk-fed infant digestive fluid"

### Supplemental Figure 1

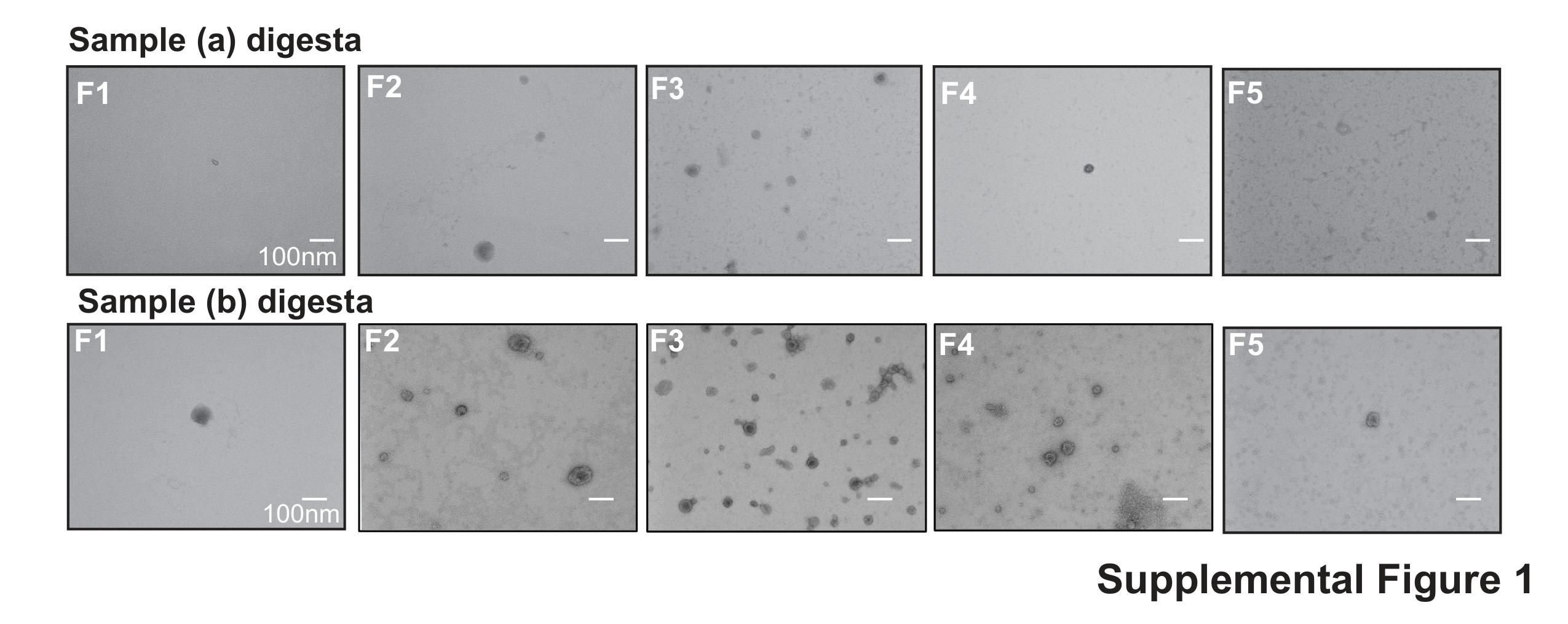
